# The N-terminus of the *Chlamydia trachomatis* effector Tarp engages the host Hippo pathway

**DOI:** 10.1101/2024.09.12.612603

**Authors:** George F Aranjuez, Om Patel, Dev Patel, Travis J Jewett

## Abstract

*Chlamydia trachomatis* is an obligate, intracellular Gram-negative bacteria and the leading bacterial STI in the United States. *Chlamydia*’s developmental cycle involves host cell entry, replication within a parasitophorous vacuole called an inclusion, and induction of host cell lysis to release new infectious particles. During development, *Chlamydia* manipulates the host cell biology using various secreted bacterial effectors. The early effector Tarp is important for *Chlamydia* entry via its well-characterized C-terminal region which can polymerize and bundle F-actin. In contrast, not much is known about the function of Tarp’s N-terminus (N-Tarp), though this N-terminal region is present in many *Chlamydia* species. To address this, we use *Drosophila melanogaster* as an in vivo cell biology platform to study N-Tarp-host interactions. *Drosophila* development is well-characterized such that developmental phenotypes can be traced back to the perturbed molecular pathway. Transgenic expression of N-Tarp in *Drosophila* tissues results in phenotypes consistent with altered host Hippo signaling. The Salvador-Warts-Hippo pathway is a conserved signaling cascade that regulates host cell proliferation and survival during normal animal development. We studied N-Tarp function in larval imaginal wing discs, which are sensitive to perturbations in Hippo signaling. N-Tarp causes wing disc overgrowth and a concomitant increase in adult wing size, phenocopying overexpression of the Hippo co-activator Yorkie. N-Tarp also causes upregulation of Hippo target genes. Last, N-Tarp-induced phenotypes can be rescued by reducing the levels of Yorkie, or the Hippo target genes *CycE* and *Diap1*. Thus, we provide the first evidence that the N-terminal region of the *Chlamydia* effector Tarp is sufficient to alter host Hippo signaling and acts upstream of the co-activator Yorkie. *Chlamydia* alters host cell apoptosis during infection, though the exact mechanism remains unknown. Our findings implicate the N-terminal region of Tarp as a way to manipulate the host Hippo signaling pathway, which directly influences cell survival.

**Author Summary:** *Chlamydia*-infected cells are known to be resistant to apoptotic cues, facilitating the successful completion of its infectious cycle. The exact molecular mechanism of apoptosis inhibition is unknown and the search for secreted effectors that mediate this is ongoing. We developed *Drosophila melanogaster* as a platform to study *Chlamydia* effector function in vivo without the confounding influence of actual infection. Genetic tools make it easy to generate transgenic flies and express bacterial effectors of interest in any tissue, allowing for discovery of new function based on observed developmental phenotypes. Using this platform, we showed that the N-terminal region of the early effector Tarp intersects with the host Hippo pathway, causing upregulation of Hippo target genes. Interestingly, the Hippo pathway directly controls the expression of potent inhibitors of apoptosis (IAPs). Our findings support a possible link between a secreted effector, Tarp and its N-terminal region, and the Hippo pathway to block apoptosis during infection.

## Introduction

*Chlamydia trachomatis* is a Gram-negative, obligate intracellular bacteria and is the leading cause of infection-induced blindness globally and the most commonly reported sexually transmitted infection (STI) in the United States, far ahead of other STIs such as Gonorrhea and Syphilis. Moreover, the year-over-year incidence of *Chlamydia* infections are consistently increasing [1], representing an ever-increasing public health burden. Despite its prevalence, much of the molecular strategies that *Chlamydia* employs to promote infection is not completely understood.

Host cell invasion is a crucial first step in *Chlamydia* infection. After attaching to the surface of the host cell, *Chlamydia* injects several bacterial effectors via type-III secretion system (T3SS) needles [2,3]. These ‘early effectors’ have unique but synergizing functions that promote efficient host cell invasion. The most well-characterized early effector is Tarp (**t**ranslocated **a**ctin-**r**ecruiting **p**hosphoprotein). It has been shown, through various approaches, that Tarp is required for efficient host cell invasion [4–6].The recruitment and remodeling of host actin is a common mechanism employed by intracellular bacteria such as *Chlamydia* to gain entry into the host cell [7–9]. In vitro, Tarp has been shown to promote the formation of F-actin as well as bundle existing F-actin filaments via different functional domains on the C-terminus[10,11]. Other early effectors such as TmeA complements Tarp function by promoting actin filament formation via the host Arp2/3 complex [12,13] while the early effector TmeB acts to finetune TmeA activity [14].

Tarp is a ∼1000 amino acid protein with distinct N- and C-termini. Tarp’s actin-remodeling domains are found on its C-terminus while the N-termini is largely devoid of annotated functional domains. Interestingly, this seemingly unassuming N-terminal configuration is observed in Tarp protein from multiple *Chlamydia* species that target a variety of host organisms [15]. For *C. trachomatis*, the N-terminus of Tarp (referred to as N-Tarp in this work) contains tyrosine-rich repeats that are rapidly phosphorylated upon delivery into the host cell [8,16,17]. The functional contribution of N-Tarp during *Chlamydia* infection is still not clearly understood though tyrosine phosphorylation is a potent post-translational modification that can engage a wide variety of signaling pathways.

We recently developed a new way to investigate effector function in vivo, using the model organism *Drosophila melanogaster* as a cell biology platform. Via transgenic expression of *Chlamydia* effectors in flies, we are able study effector function in isolation, away from the cofounding effects of active infection. Furthermore, genetic tools in *Drosophila* research allow for targeted expression of effectors in the cell or tissue of interest. Last, taking advantage of the well-understood developmental biology of *Drosophila*, we can infer effector function from the phenotypes caused by effector expression in fly tissues. We have successfully used this platform to verify that Tarp’s F-actin bundling activity also occurs in vivo and that Tarp can displace endogenous F-actin bundlers [18].

We then employed this platform to discover novel N-Tarp function in vivo in an unbiased approach. We discovered that transgenic expression of N-Tarp in flies displayed phenotypes consistent with disruption of Hippo signaling pathway [19,20]. First described in *Drosophila*, the Hippo pathway is a highly conserved signaling pathway that controls cell proliferation and cell survival, ensuring proper organ size control during development [21]. The core signaling pathway is a kinase cascade that controls the nuclear localization of the transcription co-activator Yorkie (YAP/TAZ in mammals), which, together with the DNA-binding protein Scalloped (TEAD in mammals), acts as a transcription factor to drive downstream target genes such as *Cyclin E* and *Diap1* (*XIAP* in mammals) that regulate the cell cycle and apoptosis, respectively (diagram in Fig 5). Originally discovered and characterized in *Drosophila*, perturbations in Hippo signaling lead to overgrowth or undergrowth of organs or even whole organisms [22]. Outside of its role in animal development, the Hippo pathway is also involved in wound healing and underlies several types of cancer [23,24]. In the context of *Chlamydia* infection, we indeed found evidence of changes in Hippo pathway during infection and that these changes occur in a Tarp-dependent manner [19].

In this study, we use *Drosophila* wing development as a model to study the interplay between N-Tarp and the Hippo pathway in vivo. N-Tarp expression in the developing wing leads to an increase in size, consistent with altered Hippo signaling. We use anatomical, molecular, and genetic approaches to establish that N-Tarp acts through the co-activator Yorkie to upregulate Hippo target genes. This work attributes a new function to the poorly understood N-terminal region of the early effector Tarp as a means for *Chlamydia* to manipulate the host Hippo signaling.

## Materials and Methods

### *Drosophila* stocks, handling, and rearing

From the genomic DNA of *Chlamydia trachomatis* (*Ct*) serovar L2 strain 434/Bu (ATCC VR-902B), the open reading frame that encodes for the N-terminal region of *Ct* L2 Tarp (N-Tarp, amino acid 1-431) was cloned into the pUAST *Drosophila* transformation and expression vector and used for P-element-mediated germline transformation (Model System Injections) to generate UAS-N-Tarp transgenic flies [25].

Transgene expression was performed using the modular GAL4/UAS binary expression system (ref Brand and Perrimon 1993). Briefly, the yeast-derived GAL4 transcription factor binds to its cognate UAS promoter sequence, driving transgene expression. In practice, UAS-transgene flies are crossed to select GAL4 flies to generate progeny that express the transgene of interest in the target tissues.

We previously generated and validated the transgenic expression of N-Tarp [19], as well as full length Tarp and the early effector TmeA [18] in flies.

Other stocks used were derived from or directly obtained from the Bloomington *Drosophila* Stock Center (BL) or were kind gifts from other scientists: *ban*-GFP; *nub*- GAL4 (*ban*-GFP is a gift from I. Hariharan), *ex*-lacZ/CyO;TM2/TM6B (BL-44248), *nub*- GAL4, UAS-GFP (*nub*-GAL4 derived from BL-86108), *nub*-GAL4, UAS-N-Tarp; TM6 tubGAL80 / +, *ptc*-GAL4 (BL-2017), UAS-yki:GFP (BL-28815), UAS *Diap1* RNAi HMS00752 (BL-33597), UAS *CycE* RNAi HMS00060 (BL-33654), UAS *CycE* RNAi JF02473 (BL-29314), w1118 (BL-5905), yki[B5]/CyO (BL-36290).

The cross to generate *ptc*-GAL4/UAS N-Tarp flies for compartmentalized N-Tarp expression in the adult wing was reared at 18°C to improve progeny survival. All fly stocks and other crosses were reared at 25°C. Flies were grown on Nutri-fly BF media (Genesee Scientific) supplemented with 0.45% v/v propionic acid as mold inhibitor. We used FlyBase (release FB2024_02) to obtain information on phenotypes, expression pattern, and available stocks [26].

### Immunostaining and imaging of imaginal wing discs

Wing imaginal discs were dissected from third instar larvae in PBS and transferred into a watch glass well. Tissue fixation was performed using 4% formaldehyde in PBS-triton-x-100 (0.1% v/v)(PBT) for 10 minutes while rocking on a small orbital shaker. Fixed tissue were blocked in PBT-BSA for 1 hour prior to immunostaining. Primary antibodies used: 1:50 anti-LacZ (40-1a, DSHB). Secondary antibodies used: 1:400 anti-mouse Alexa Fluor 594 (Invitrogen). Co-stains used: 1:400 phalloidin Alexa Fluor 568 (Invitrogen), 1:400 GFP-booster (Chromotek), DAPI (Invitrogen). Stained wing discs were mounted onto glass slides using Aqua-polymount and placed under a cover glass (#1.5).

A Zeiss LSM 710 confocal microscope was used to acquire fluorescent images of immunostained wing discs using a 10x EC Plan-Neofluar objective. Imaging of *ban*-GFP and *ex*-LacZ reporter intensities were performed with identical acquisition settings between control and experimental samples.

### Measuring wing pouch size

GFP expression driven by *nub*-GAL4 visually delineates the wing pouch region. Z-stacks of confocal images of wing discs were z-projected using the ‘maximum intensity’ setting. The region of interest defined by GFP expression was selected and the ROI area measured. Image analysis was performed using FIJI [27].

### Scoring and imaging of wing phenotypes in adult flies

Adult fly wings were visually examined under CO_2_ anesthesia using a Leica stereomicroscope and categorized and counted as having crumpled, wavy, or straight wings (representative images in Fig 4). Statistical tests were performed per phenotypic category between genotypes. Comparisons between two genotypes (as in Fig 4B,D) used paired t test, while comparison across three genotypes (as in Fig 4E) used Browne-Forsythe and Welch ANOVA tests with Dunnett’s T3 multiple comparisons.

Flies were mounted on wooden picks for imaging attached wings. To analyze wing dimensions, wings were clipped at the hinge, fixed in 70% ethanol, and mounted on glass slides with a small amount of 40% glycerol. Whole flies or clipped wings were imaged using either a Zeiss Stemi 508 stereomicroscope with an Axiocam 208 color camera or a Keyence VHX-7000 digital microscope. Measurement of wing dimensions were performed using FIJI [27].

For quantifying trichome number in the wing, a defined square ROI was overlayed within the *ptc*-GAL4 wing compartment, close to the wing margin (see Fig S3) and the number of trichomes within the ROI was counted visually.

### Measuring Hippo pathway activity using genetically encoded reporters

To measure the Hippo pathway activity, we utilized two genetically encoded Hippo pathway reporters: *ban*-GFP [28] and *ex*-LacZ [29]. N-Tarp was expressed in the wing disc using *nub*-GAL4 in the genetic background of either *ban*-GFP or *ex*-LacZ. The wing discs were dissected and immunostained for either GFP (using GFP-booster) or LacZ and imaged using confocal microscopy (see methods above). Z-stacks were z-projected using the ‘sum slices’ setting to generate a cumulative image of reporter intensity through the entire wing disc thickness. For *ban*-GFP, the pixel intensity was recorded along a line ROI placed on the wing disc midline. For *ex*-LacZ, the mean pixel intensity within an area ROI delineating the wing pouch region was measured. Image analysis was performed using FIJI [27].

### In vitro kinase assay of Purified N-Tarp

GST-tagged N-Tarp (GST-Tarp^1-625^) was expressed in *E. coli* and purified using glutathione affinity purification as previously described [11]. *Drosophila* lysate was obtained by mechanical disruption of frozen flies with mini mortars and pestles followed by sonication in 100mM KCl, 2mM MgCl2, 1mM ATP and 10mM HEPES (pH 7.4) (Buffer A). Insoluble *Drosophila* material was cleared by centrifugation (20,000 X *g*, 20 min, 4°C). Purified N-Tarp and a GST control immobilized on glutathione beads were incubated with *Drosophila* lysate for 1 hour at 20°C. Following the incubation, beads were washed 3X with buffer A and resuspended directly into Laemmli sample buffer. Samples were analyzed by SDS-PAGE followed by Coomassie stain or Western blot for tyrosine phosphorylation (4G10 mouse monoclonal antibody, EMD Millipore Corp.).

### Graphs, Figures, and Statistics

Graphs and statistical analyses were generated and performed using Graphpad Prism 10. Representative images and schematics were prepared using FIJI, Adobe Photoshop, and BioRender. Figures were assembled in Adobe Illustrator.

## Results

### Expressing N-Tarp in the wing imaginal disc results in tissue overgrowth

Imaginal discs are larval tissue that serve as precursors to major structures in the adult fly, such as the antenna, eye, legs, and wings, among others. As the larva grows, the imaginal discs such as the wing also proportionally increase in size [30]. This growth is driven by simultaneous cell proliferation and tissue patterning mediated via the Hippo pathway [31], making the wing imaginal disc an excellent tissue to study Hippo signaling. The pair of larval wing discs form the complete dorsal thorax and wings. The wing pouch region of the larval wing disc (Fig 1A, middle) gives rise to the adult wing blade (Fig 1A, right).

**Figure 1.**
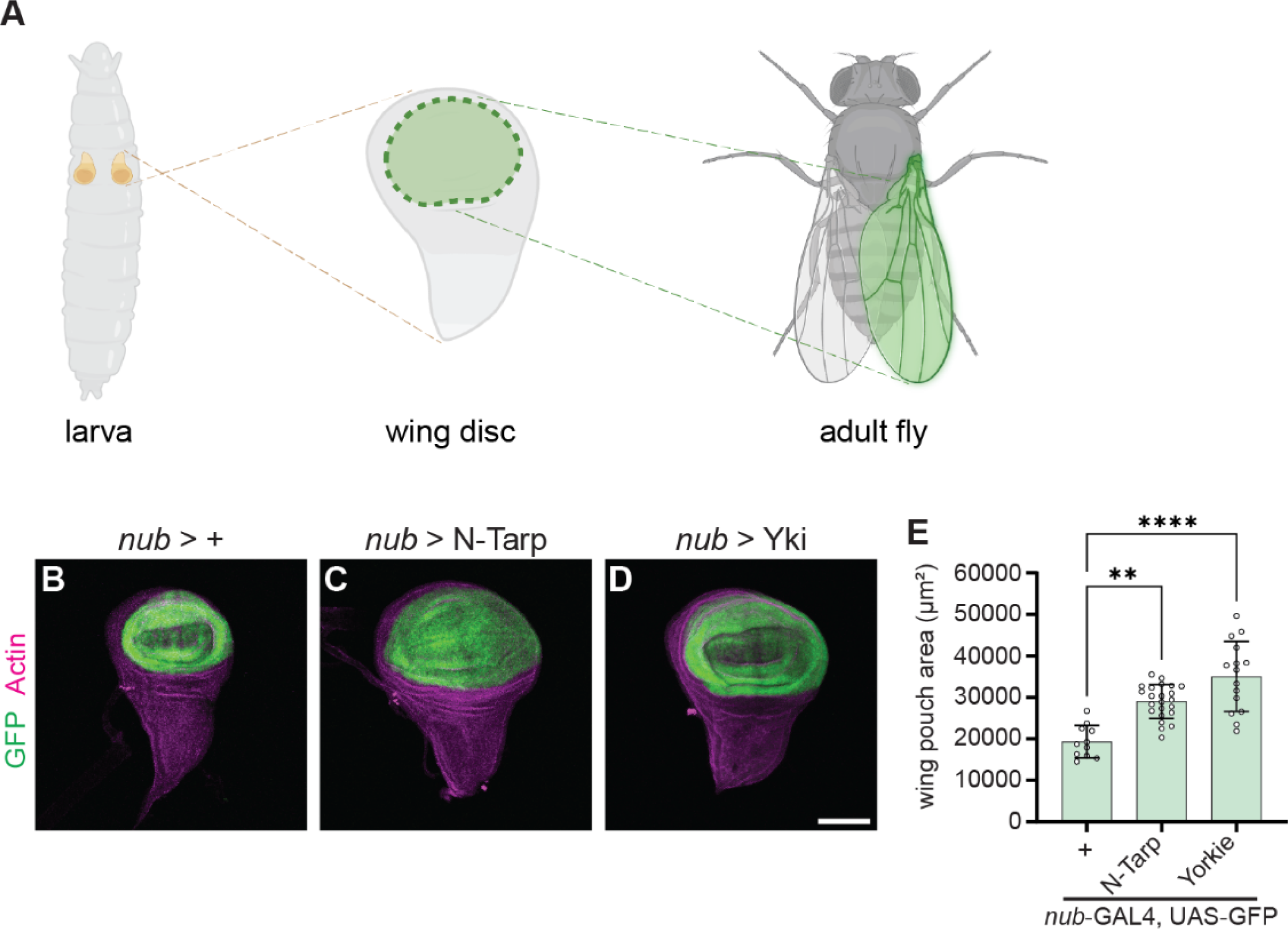
N-Tarp expression causes larval wing pouch overgrowth. (A) *Drosophila* larvae carry a pair of imaginal wing discs (left). The wing discs are larval tissue precursors of the adult dorsal thorax and wings. The pouch region of the wing disc (middle, green region) give rise to the entire wing blade of the adult fly (right, green highlight). Created in BioRender. (B-D) Wing discs were dissected from 3^rd^ instar larvae and co-stained with phalloidin to label actin. *nub*-GAL4 drives expression of (B) GFP alone, or together with (C) N-Tarp or (D) the Hippo pathway co-activator Yorkie (Yki). Scale bar represents 50µm. (E) Wing pouch area measurement from control discs or discs expressing N-Tarp or Yorkie. Each hollow dot represents a wing disc. Bar graphs indicate the mean and SD. Kruskal-Wallis test with Dunn’s multiple comparisons was used (**, p<0.01; ****, p<0.0001).

We used *nubbin*-GAL4 (*nub*-GAL4) to express N-Tarp in the wing pouch region (Fig 1B, GFP-expressing region). By measuring the area of GFP-positive tissue, we show that N-Tarp expression results in a larger wing pouch area compared to control (Fig 1C, E). Overexpression of Yorkie, the Hippo pathway transcription co-activator, similarly results in increased wing pouch area (Fig 1D, E), consistent with published findings [32,33].

We then allowed *nub*>N-Tarp larvae to develop into adult flies to look at impact on the adult wing blade. The wings of control adults are flat when fully expanded (Fig 2A). Surprisingly, the vast majority (94.9%, n=350 flies over 3 trials) of *nub*>N-Tarp adult flies have crumpled, unexpanded wings (Fig 2B). The crumpled wings remain throughout the life of the adult flies. Counterintuitively, wing overgrowth has been observed to present as crumpled, unexpanded wings [33,34]. Indeed, the small proportion (5.1%) of *nub*>N-Tarp adult flies that have fully expanded wings display a much larger wing area (Fig 2D) compared to control flies (Fig 2C), consistent with the wing pouch overgrowth seen in *nub*>N-Tarp larval wing discs (Fig 1). As expected, overexpression of Yki in the wing pouch also causes increased wing size (Fig 2E). Interestingly, this increased wing size was specific to N-Tarp expression and was not observed upon expression of full-length Tarp or another *C. trachomatis* early effector, TmeA (Fig S1).

**Figure 2.**
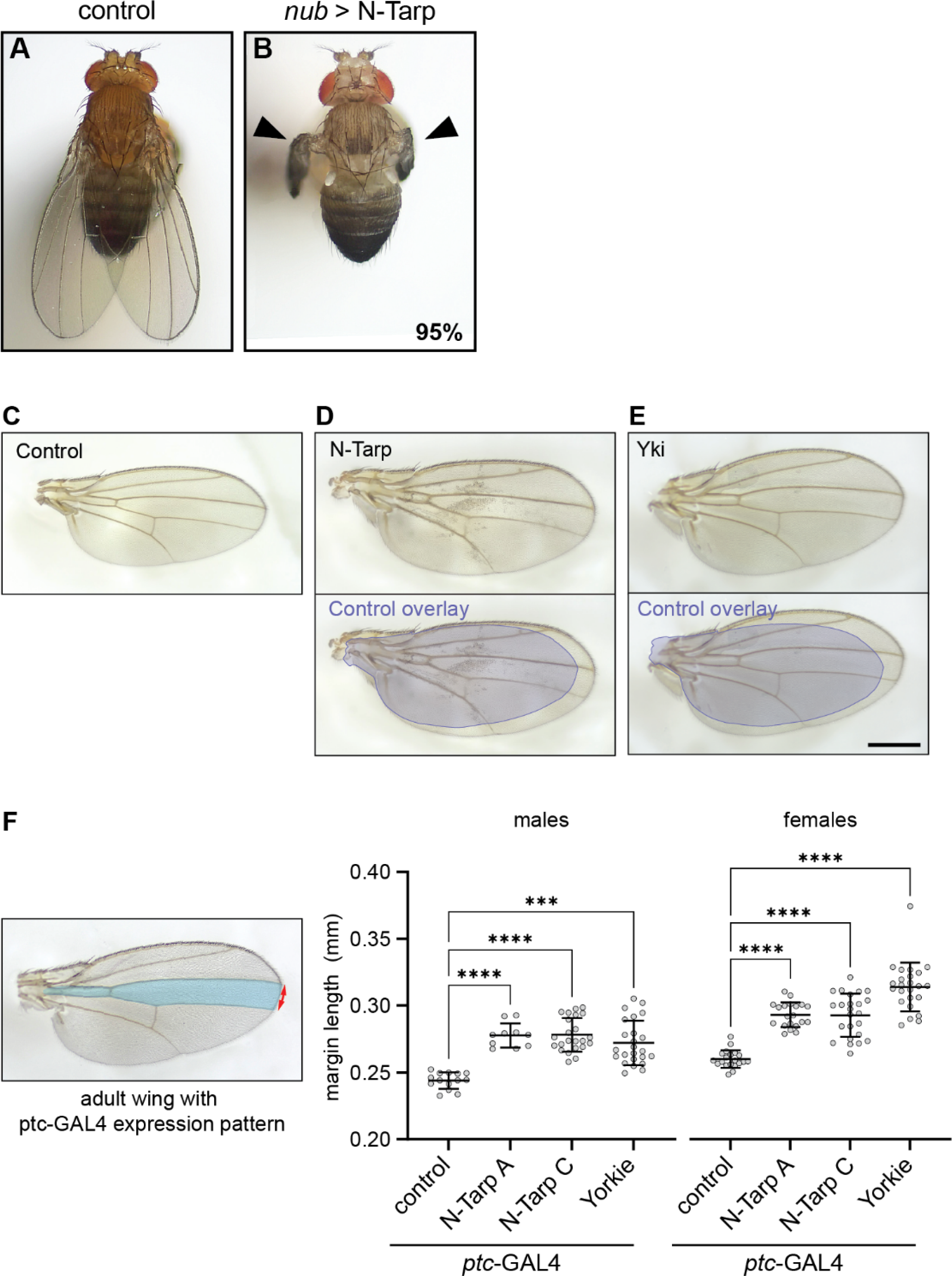
N-Tarp expression leads to increased wing size. (A-B) Dorsal view of an adult (A) control (*nub*-GAL4/+) fly or (B) *nub*-GAL4/UAS-N-Tarp fly. Black arrowheads indicate crumpled, unexpanded wings that occur in 95% of examined flies (n=350 flies over 3 trials). (C-E) Dissected wing blades were mounted on slides and imaged. (C) Control wings are flat with stereotypical vein and crossvein patterns. (D) 5% of *nub*-GAL4/UAS-N-Tarp flies have fully expanded wings (top panel). Bottom panel is the same *nub*-GAL4/UAS-N-Tarp wing with an overlay of the control wing outline (blue), revealing the difference in wing size. (E) Wing from *nub*-GAL4/UAS-Yorkie (Yki) flies are also larger compared to control. Scale bar is 500µm. (F) (left) *ptc*-GAL4 expression pattern (blue region) is limited to a small compartment of the adult wing. The distance between the top and bottom boundary of the expression pattern was measured along the wing margin (red arrows). (right) Scatter plot of wing margin length along the *ptc*- GAL4 expression boundary in control, UAS-N-Tarp, and UAS-Yorkie wings, indicating mean and SD. Two unique transgenic lines of UAS-N-Tarp (A & C) were tested, showing similar increase in wing margin length. Yorkie expression also results in increased wing margin length. Wings from male and female adult flies were analyzed separately. Kruskal-Wallis test with Dunn’s multiple comparisons was used (***, p<0.001; ****, p<0.0001).

To increase the number of analyzable, fully expanded wings, we limited the expression of N-Tarp to a small compartment of the adult wing along the anterior/posterior boundary using the *ptc*-GAL4 driver (Fig 2F, blue region in left panel) [35]. Indeed, N-Tarp expression in a limited region of the wing did not interfere with wing expansion. We then measured the wing margin length between the anterior and posterior boundaries of the *ptc*-GAL4 domain (Fig 2F, red arrow in left panel). Multiple, unique N-Tarp transgenic lines both resulted in increased margin length compared to control wings (Fig 2F, graph). Moreover, This N-Tarp-induced increase in margin length phenocopies the increase caused by Yorkie overexpression (Figure 2F), consistent with the observations from the larval wing disc.

A larger tissue size can arise from an increase in the total cell number from overproliferation, consistent with the role of the Hippo pathway. It can also occur from an increase in the individual cell size with no appreciable increase in cell number, as in the case of TOR signaling [36]. To distinguish between the two possibilities, we quantified the cell density within the *ptc*-GAL4 compartment upon expression of N-Tarp or Yorkie overexpression by counting the number of trichomes, microscopic bristles that cover the entire wing surface, within a defined region [37]. Trichomes originate from individual cells and act as a surrogate visual indicator of cell number. We observed no change in trichome density between control, N-Tarp-expressing, and Yorkie-overexpressing wings (Fig S2), indicating that larger wing dimensions upon N-Tarp expression is likely due to increased cell number as a result of overproliferation.

### N-Tarp causes upregulation of Hippo pathway target genes

Above, we show that N-Tarp-induced tissue overgrowth in the larval wing disc and in the adult wing phenocopies changes brought on by Yorkie overexpression. To further support the link between N-Tarp and the Hippo pathway, we sought to determine whether the expression of canonical Hippo target genes is also altered in the presence of N-Tarp. The expression of the gene *bantam* is controlled by the Hippo pathway [38,39]. We used a *bantam*:GFP reporter [28] to query Hippo pathway activity in developing wing discs upon expression of N-Tarp. Control wing discs display an overall low GFP fluorescence in the wing pouch, with only very small regions of moderate activity (Fig 3A). On the other hand, N-Tarp expression in the wing pouch resulted in a striking increase in *bantam* gene expression, as evidenced by the expanded area of moderate GFP fluorescence intensity and the appearance of high activity regions (Fig 3B). Quantitative analysis of GFP fluorescence intensity along a linear ROI (Fig 3A,B, dashed line) shows a consistently increased level of *bantam* gene expression upon N-Tarp expression over multiple wing discs examined (Fig 3C). Instead of measuring GFP intensity over the wing pouch region, a linear ROI was chosen to remain independent of the different wing pouch sizes between the two genotypes (Fig 1E). A transgenic reporter of another canonical Hippo pathway target gene, *expanded*, [40] also showed similar results (Fig 3D-F). In all, the above results provide molecular proof that N-Tarp expression alters Hippo pathway activity.

**Figure 3.**
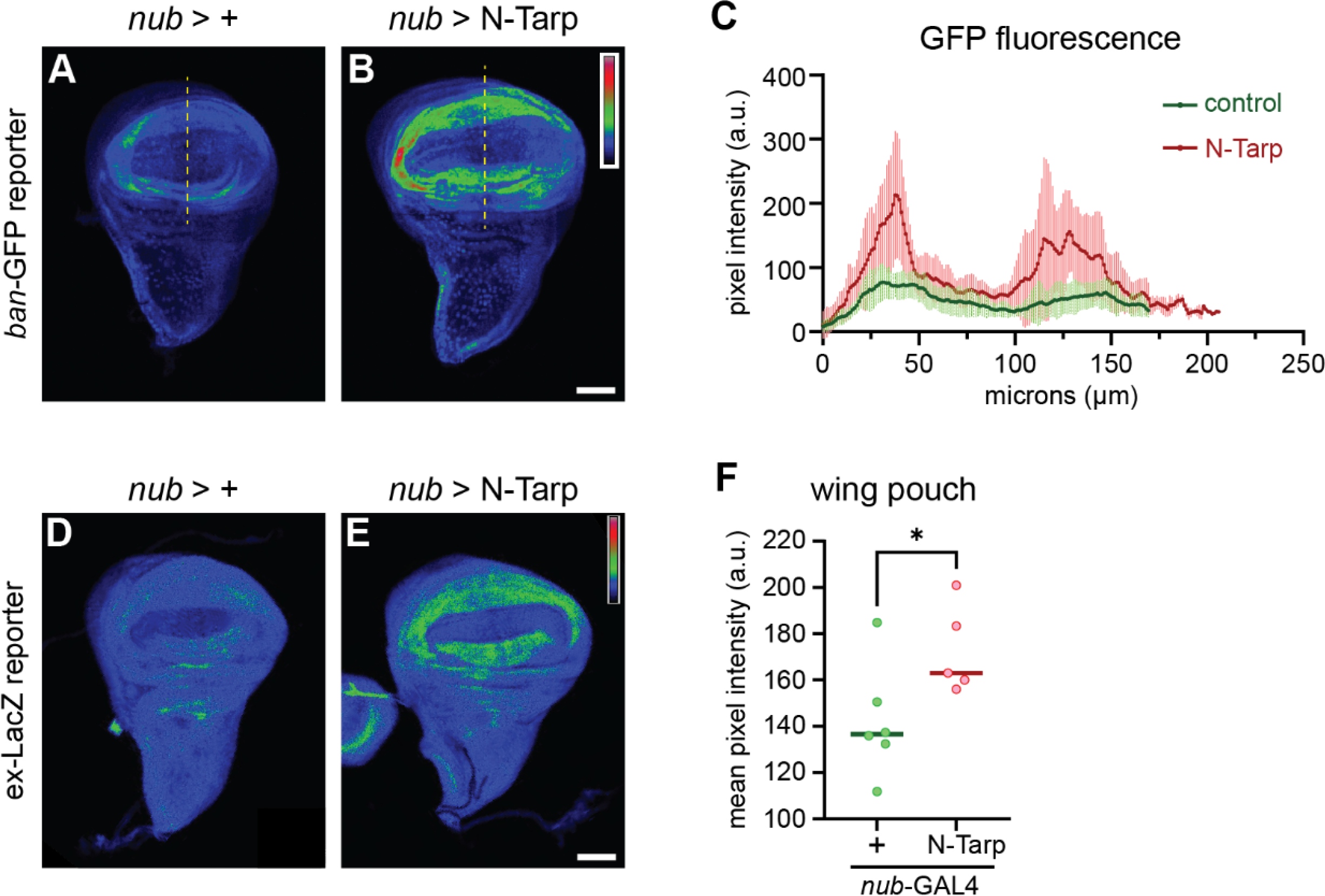
Hippo pathway target genes are upregulated upon N-Tarp expression. (A-C) The promoter of the Hippo pathway target gene *bantam* (*ban*) controls the expression of GFP (*ban*-GFP) and used as a reporter of Hippo pathway activity in the wing discs. GFP fluorescence intensity is represented using a rainbow LUT. (A) Control (*nub*>+) wing discs display the baseline GFP intensity while (B) N-Tarp expression in the wing pouch (*nub*>N-Tarp) results in increased GFP intensity. (C) Mean pixel intensity ± SD along a line ROI (yellow dashed lines) that traverses the wing pouch were plotted for control (*nub*>+; green; n=22) and N-Tarp (*nub*>N-Tarp; red; n=17). (D-F) Another Hippo pathway reporter uses the promoter of the Hippo target gene *expanded* (*ex*) to drive LacZ expression (*ex*-LacZ). LacZ levels were visualized by immunostaining and displayed using a rainbow LUT. (D) Control (*nub*>+) wing discs show baseline LacZ levels while (E) N-Tarp expression in the wing pouch (*nub*>N-Tarp) results in elevated LacZ levels. (F) The mean pixel intensity ± SD within the wing pouch region for control (*nub*>+; green) and N-Tarp (*nub*>N-Tarp; red) is plotted. Welch’s test used (*, p<0.05). Scale bar is 50µm.

### N-tarp-induced wing overgrowth is rescued by disrupting the Hippo pathway

To prove that N-Tarp function acts through the Hippo pathway, we attempted to rescue the crumpled wing phenotype (Fig 2B) by either disrupting: 1) the Yki transcription factor; or 2) the canonical Hippo target genes *Diap1* and *CycE*. Together, *Diap1* (*Drosophila inhibitor of apoptosis 1*) and *CycE* can promote tissue growth through overproliferation. If N-Tarp’s in vivo activity is indeed acting through the Hippo pathway, then disrupting either the key transcription factor, Yki, or the canonical Hippo target genes linked to cell proliferation, should result in amelioration of the crumpled wing phenotype.

We expressed N-Tarp in the wing pouch of *yki*^+/-^ heterozygous null flies. Reduced *yki* gene dosage leads to reduced Yki protein levels [41] without dramatically affecting animal viability—the use of RNAi to knockdown expression of *yki* in the wing pouch leads to strong lethality (data not shown). We then measured the frequency of observing N-Tarp-induced crumpled wings in the wildtype background vs *yki*^+/-^ background. As expected, expression of N-Tarp in the wildtype background results in all flies with abnormal wings with a high frequency of crumpled wings (Fig 4A, left panel; 4B). Expression of N-Tarp in the *yki*^+/-^ background results in reduced crumpled wing frequency, an increase in an intermediate, wavy wing phenotype frequency, and, more importantly, the appearance of normal, straight wings (Fig 4A, middle and right panels; 4B). We also expressed N-Tarp in the wing pouch while simultaneously knocking down *Diap1* or *CycE* expression by RNAi. Knockdown of either *Diap1* or *CycE* shifts the distribution of N-Tarp-induced wing phenotypes more strongly towards the intermediate wavy wing or fully straight wings (Fig 4C-E). Flies that that are solely heterozygous null for Yki or underwent RNAi knockdown of *Diap1* or *CycE* in the wing pouch all have straight wings (Fig S3).

**Figure 4.**
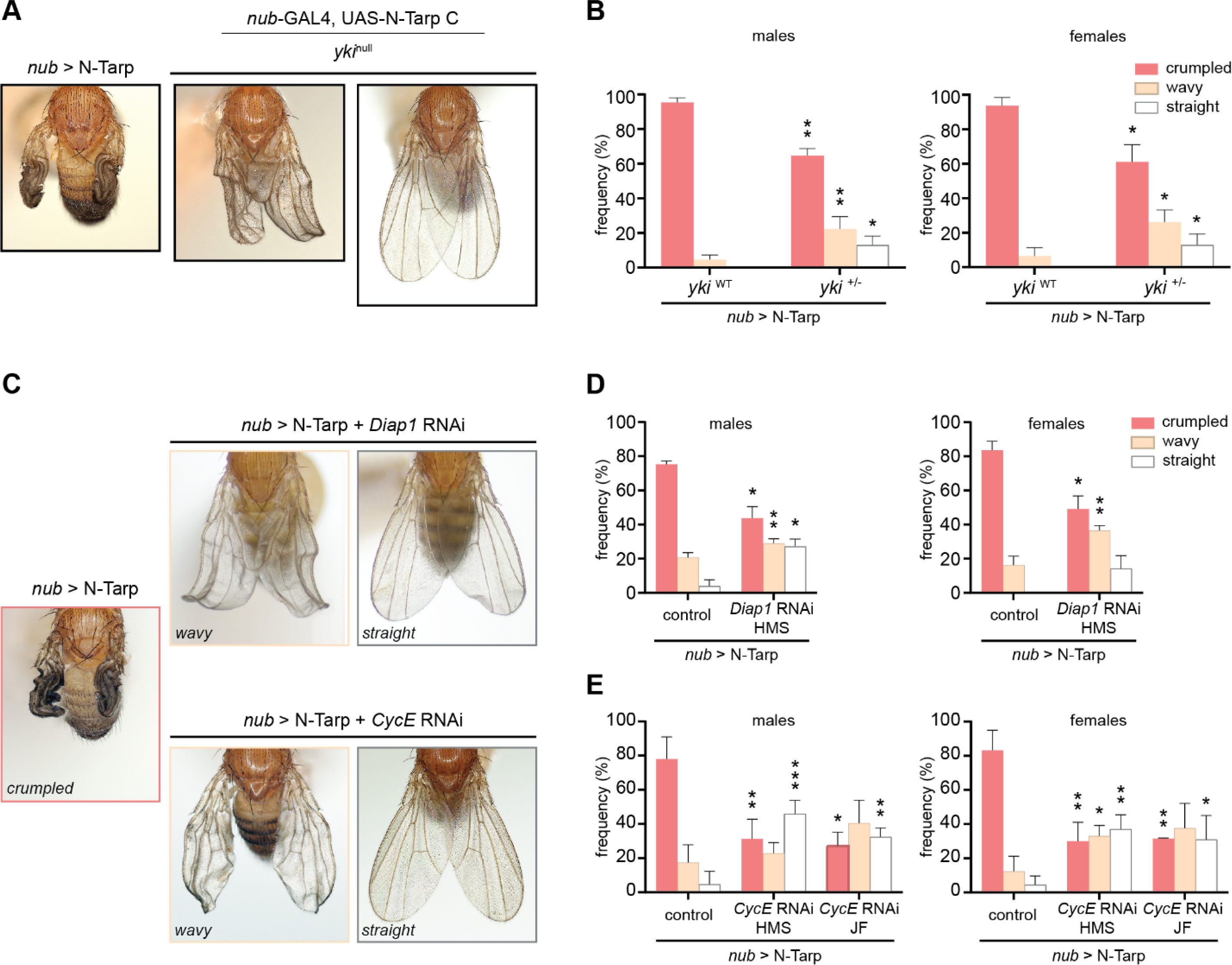
N-Tarp-induced overgrowth rescued by disrupting the Hippo pathway. N-Tarp was expressed in the developing wing on its own (*nub*>N-Tarp) or in the genetic background of (A-B) heterozygote yki^null^ mutant; or (C-E) RNAi knockdown of the Hippo target genes *Diap1* or *CycE*. For each condition, the severity of wing phenotypes was categorized and scored as crumpled (strong), wavy (intermediate), or straight (normal). Representative images of wing phenotypes are displayed (A,C). Expression of N-Tarp in the developing wing results in a crumpled wing phenotype in 77%-95% of the progeny (B, yki^WT^; D-E, control). When expressed in heterozygous yki^null^ background, there is a significant decrease in the proportion of flies displaying crumpled wings and a concomitant increase in the flies displaying an intermediate wavy wing phenotype and the appearance of flies with normal wings (B, yki^+/-^). Expression of N-Tarp in a *Diap1* RNAi or *CycE* RNAi background similarly caused a drop in crumpled wing frequency and a shift towards the intermediate wavy and normal wing category (D,E). HMS and JF refer to different *Drosophila* RNAi collections. Statistical tests were performed per phenotypic category, more details in methods (*, p<0.05; **, p<0.01; ***, p<0.001). Four independent trials of the genetic rescue experiments were performed.

A shift from predominantly crumpled wings to partially or fully expanded wings upon disruption of the Hippo pathway, as performed above, strongly indicates that N-Tarp exerts its function in vivo by acting through the Hippo pathway. The essential nature of Hippo pathway activity to animal viability precludes the examination of N-Tarp function in a completely null Yki, *Diap1*, or *CycE* background, which can explain the partial nature of the genetic rescue.

In summary, using *Drosophila* cell and developmental biology as a platform, we present strong evidence that the N-terminal region of Tarp alters the activity of the host Hippo signaling pathway upstream of the co-activator Yorkie, resulting in the increased expression of Hippo target genes (Fig 5). This agrees with our previous study that shows a Tarp-dependent change in Hippo pathway activity in an in vitro infection model [19]. Finally, this work provides ample evidence for a novel function of the N-terminal region of Tarp and positions Tarp as having an important role outside of host cell invasion.

**Figure 5.**
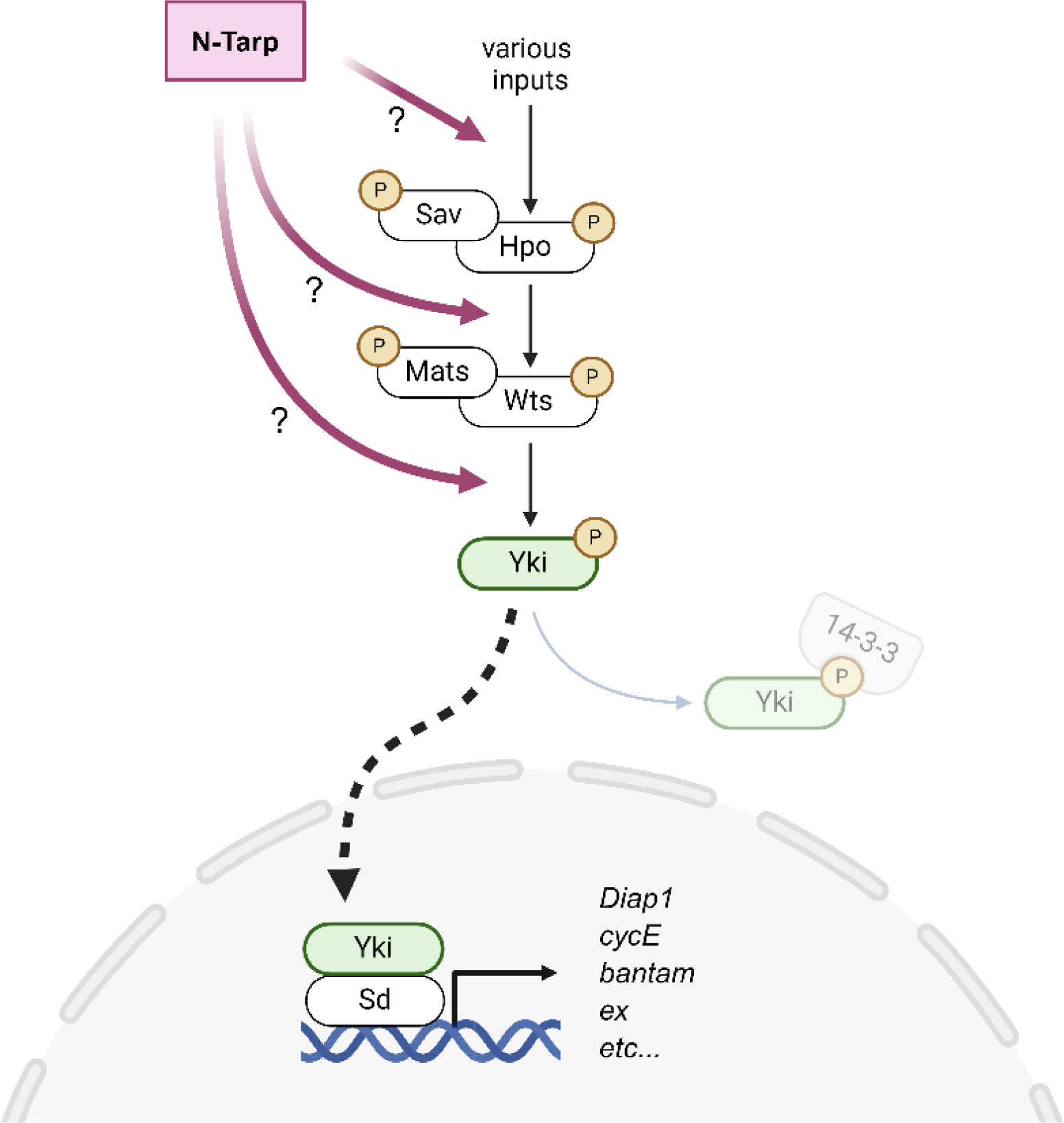
The N-terminal region of the *Chlamydia* effector Tarp acts on the host Hippo pathway to increase target gene expression. The core kinase cascade of the host Hippo pathway is made up of the following proteins: Salvador (Sav), Hippo (Hpo), Mob as tumor suppressor (Mats), and Warts (Wts). The Hippo pathway incorporates multiple signaling inputs to set the phosphorylation state of the transcriptional co-activator Yorkie (Yki). Phosphorylated Yki binds to 14-3-3 protein and retained in the cytoplasm. Unphosphorylated Yki translocates into the nucleus and binds to the TEAD protein Scalloped (Sd), which then acts as a transcription factor to turn on Hippo target gene expression. N-Tarp acts upstream of the Yki co-activator to cause increased expression of canonical Hippo target genes but whether it acts directly on the core signaling components or as an upstream input is not yet determined. Created in BioRender.

## Discussion

This work takes advantage of the genetic tools and techniques available to *Drosophila* researchers to characterize the in vivo function of the N-terminal region of Tarp using morphological, molecular, and genetic approaches. In particular, demonstration of genetic rescue of the N-Tarp-induced wing phenotype conclusively shows that N-Tarp acts through the Hippo pathway, placing it upstream of the co-activator Yorkie. Regulation of the Hippo pathway is complex, with multiple upstream inputs as well as direct influence on intermediate components [42]. As such, further efforts to determine the molecular mechanism of N-Tarp activity will involve identifying the exact point of intersection with the Hippo pathway (Fig 5).

The function of the N-terminal region of Tarp is not well-understood. The early effector Tarp is secreted into the host cell to promote host cell invasion after *Chlamydia* attachment to the cell surface [6,8]. The importance of Tarp in the invasion process is highlighted by its evolutionary conservation across multiple *Chlamydia* species that target a wide spectrum of host animals, ranging from mammals, birds, reptiles, to fish [15]. The C-terminal region of Tarp harbor actin-interacting domains that is important for actin nucleation, polymerization, and filament bundling [4,10,11] whereas the N-terminal region is mostly devoid of any annotated domains. The exception to this is the human pathogen *C. trachomatis*, whose Tarp N-terminus can contain varying number of tyrosine-rich repeats, depending on the serovar [17].

The presence of a Tarp N-terminal region is observed across multiple *Chlamydia* species [15], a telltale sign of a significant but underappreciated role in infection. *C. trachomatis* Tarp is rapidly phosphorylated at the N-terminus upon delivery into the host cell, during the attachment and early invasion steps of infection [8,43]. Similarly, when expressed in *Drosophila* tissue, we observe phosphorylation of full-length Tarp [18] and the N-Tarp fragment is phosphorylated when incubated in *Drosophila* lysate (Fig S4). Phosphorylation is a powerful signaling cue, and peptides that represent N-Tarp phosphorylation have been shown to bind strongly to human SHC1, an SH2-domain-containing protein, to activate MEK/ERK signaling and drive downstream gene expression related to cell survival [44].

*Chlamydia*-infected cells in culture are surprisingly resistant to extrinsic and intrinsic triggers of apoptosis [45]. *Chlamydia* development takes 24-48 hours to complete depending on the host cell type. This is ample time for the host cell to mount a defense response and, failing that, to undergo apoptosis. Once triggered, apoptosis progresses rapidly, resulting in cell death in a shorter time frame. However, apoptosis in *Ct*-infected cells is blocked at multiple levels—there is no activation of Bcl-2 family members Bax and Bak [46,47], no release of cytochrome c from mitochondria, and lack of processing of caspases 9 and 3 [48]—though it is not clear how this is accomplished. The work by Mehlitz et al. (2010) as well as studies focused on another *C. trachomatis* effector, CpoS, both address the search for *Chlamydia*’s anti-apoptotic effectors [49].

Inhibitor of Apoptosis (IAP) proteins are negative regulators of apoptosis and cell death that can block apoptosis [50]. Interestingly, the human and *Drosophila* IAPs *XIAP* and *Diap1*, respectively, are canonical target genes of the host Hippo pathway. It was recently shown that Hippo pathway activity is altered during *Ct* infection in vitro, resulting in increased nuclear localization of the co-activator YAP [51], as well as increased *XIAP* gene expression [19]. Moreover, the increase in *XIAP* expression is dependent on the presence of Tarp [19]. Our study extends that discovery by implicating the N-terminal region of *Ct* Tarp as sufficient to alter Hippo signaling using an in vivo model. Taken together, it is possible that *Chlamydia*, via the N-terminal region of Tarp, also engages the Hippo pathway to increase IAP levels to ensure host cell survival during infection.

The Hippo pathway crosstalks with other host signaling pathways, apart from its classic role in controlling host cell survival. Therefore, it is not surprising that many other pathogens have evolved strategies to directly manipulate the host Hippo pathway to promote infection. *Legionella pneumophila* secretes the effector LegK7, which directly phosphorylates MOB1 (Mats in flies), altering the host transcriptional response to promote infection [52]. MOB1/Mats is a member of the core kinase cascade of the Hippo pathway and is normally phosphorylated by the upstream Hippo kinase. *Ehrlichia chaffeensis* secretes TRP120 which turns down Hippo signaling, leading to expression of the anti-apoptotic Hippo target gene *SLC2A1/GLUT1* [53]. Viral infections have also targeted Hippo signaling. The human papilloma virus (HPV) E6 protein directly interacts with multiple members of the Hippo pathway [54,55], contributing to the formation of the poorly differentiated tissue lesions associated with HPV infections. *Helicobacter pylori*, as well as many other viruses have documented changes in Hippo signaling, although the mediating effector or viral protein might act indirectly or has not been elucidated at this time [56,57].

This work supports a role for Tarp, via its N-terminal region, as the mediating effector that intersects with the host Hippo pathway. The engagement of Hippo signaling during *C. trachomatis* infection is a newly emerging avenue of inquiry that may hold answers to long-standing questions of resistance to apoptosis as well as pathological changes in infected tissue in vivo.

## Acknowledgements

This work was supported by grants awarded to TJ (NIH R21AI148999 and R01AI139242) and startup funds awarded to GA. The authors acknowledge the help of undergraduates Gian Viray and Rom Peles in *Drosophila* stock creation and maintenance. Stocks obtained from the Bloomington *Drosophila* Stock Center (NIH P40OD018537) were used in this study. Thanks to Dr. Sophia Friesen and the Hariharan lab (UC Berkeley) for sharing fly stocks. Monoclonal antibodies were obtained from the Developmental Studies Hybridoma Bank, created by the NICHD of the NIH and maintained at The University of Iowa.

## Supporting Information

**Figure S1.**
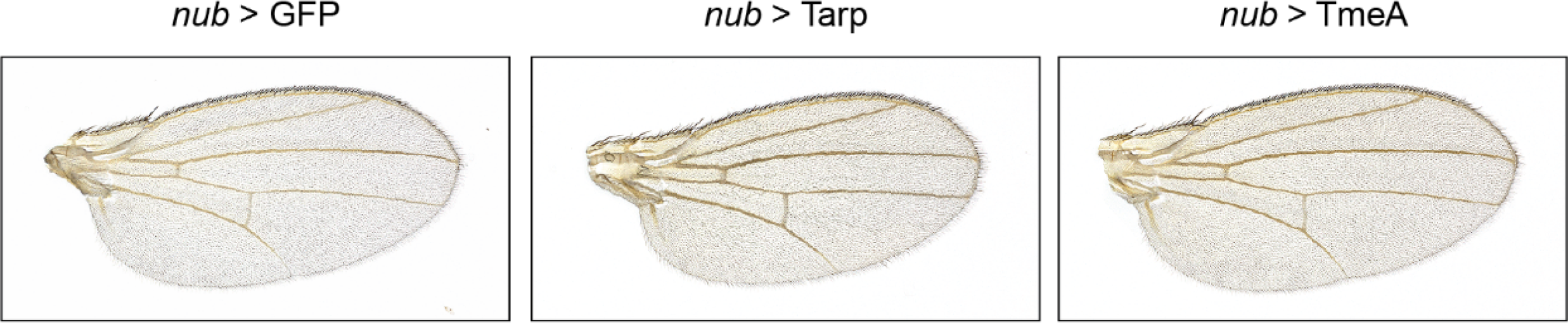
Expression of *Chlamydia* effectors Tarp (full-length) or TmeA does not disrupt wing morphology. Representative wing images from flies expressing green fluorescent protein (GFP), full-length Tarp, and another *C. trachomatis* early effector, TmeA, in the developing wing pouch using *nub*-GAL4.

**Figure S2.**
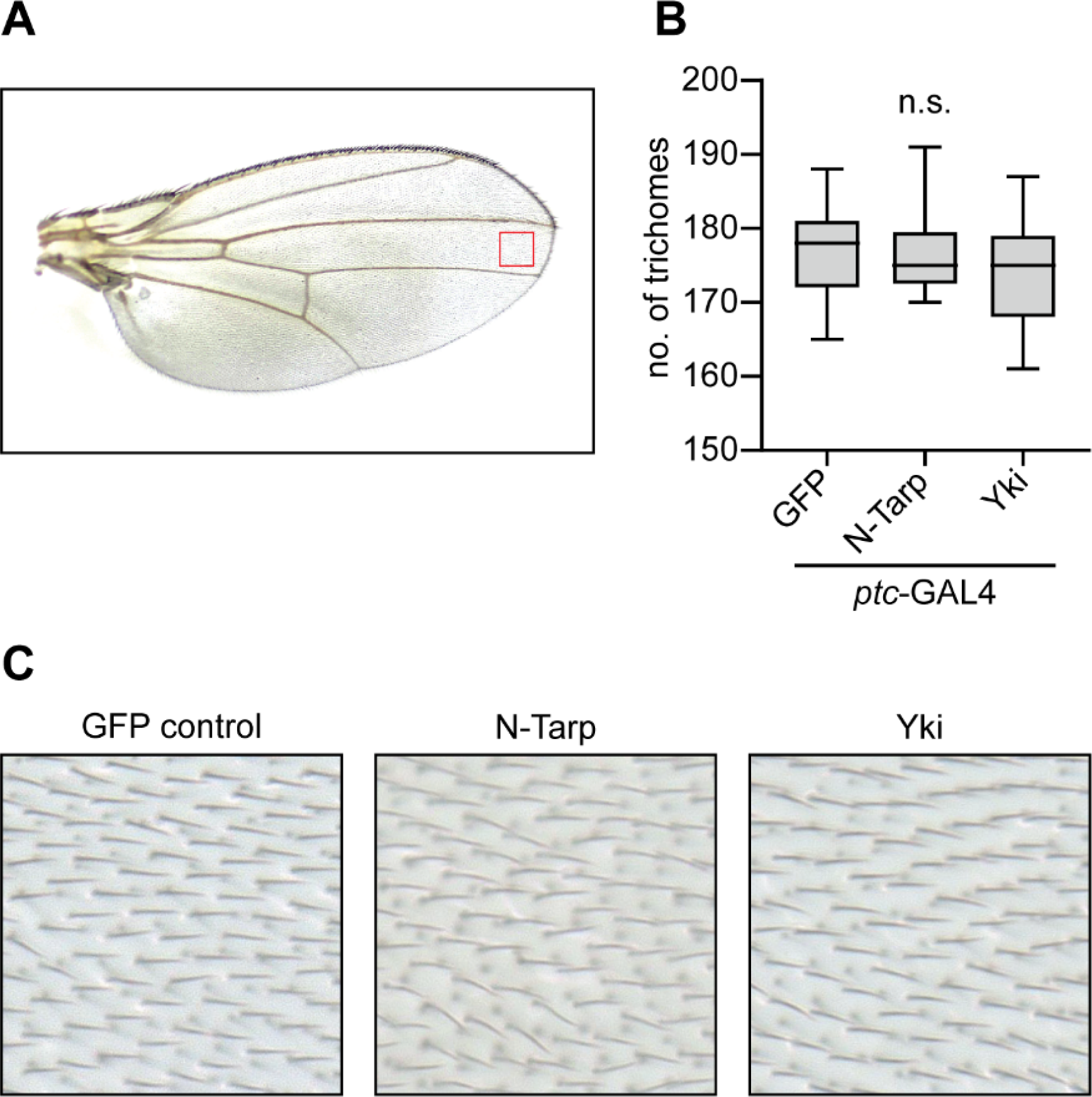
Trichome density in the wing is not affected by the expression of N-Tarp or Yorkie. (A) A representative control wing displaying the region analyzed for trichome density (red box). (B) Box and whisker plots represent the number of trichomes (microscopic wing bristles) within the defined analysis region (red box) in wings expressing GFP (negative control), N-Tarp, or Yorkie (Yki) driven by *ptc*-GAL4. There is no statistical difference (n.s.) between the groups (Kruskal-Wallis test, n≥10 per group). (C) Representative, high magnification images of trichomes within the defined analysis region from wings expressing GFP (negative control), N-Tarp, or Yorkie (Yki) driven by *ptc*-GAL4.

**Figure S3.**
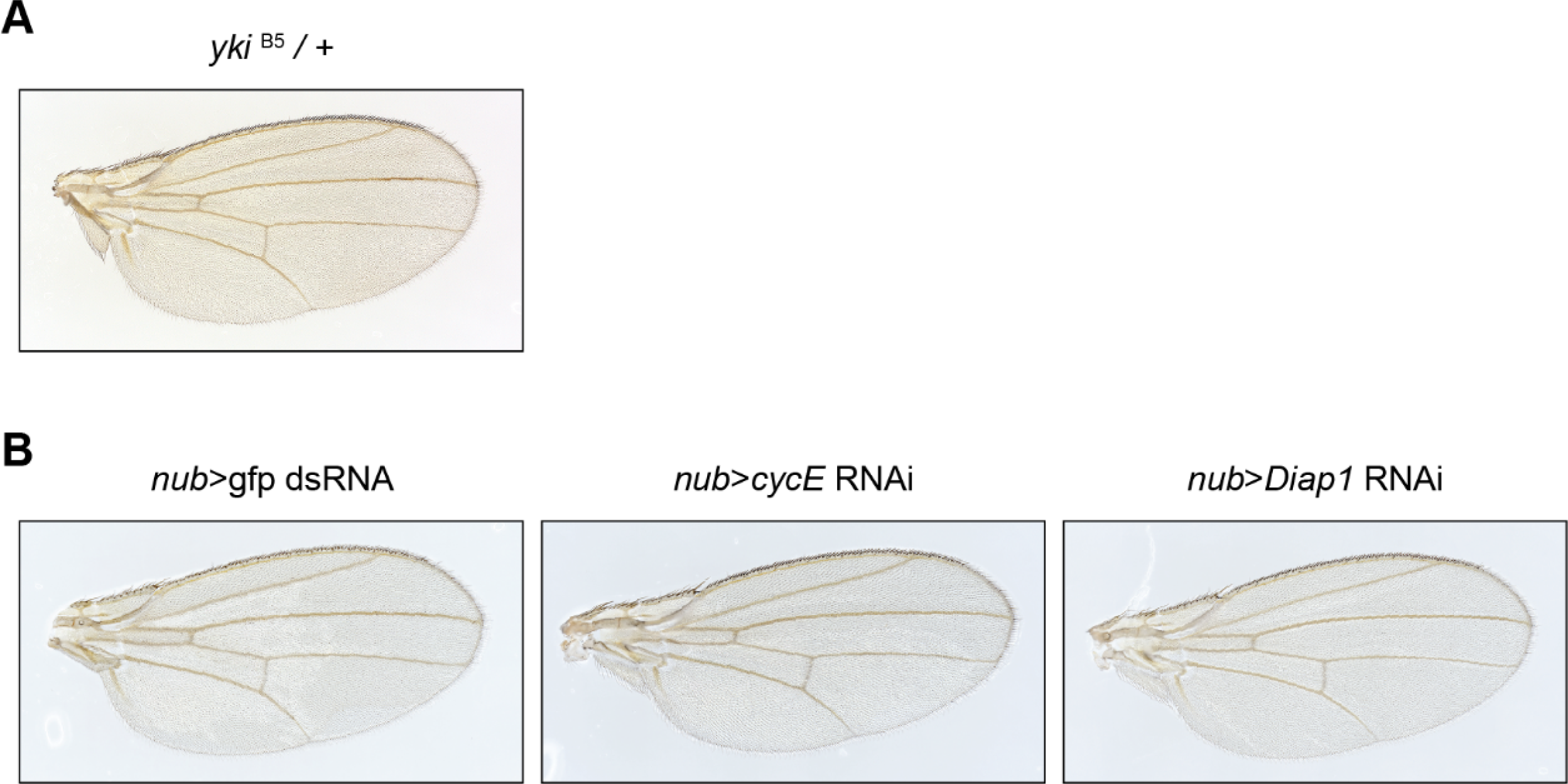
Altering the levels of Yorkie, Diap1, or CycE alone does not disrupt wing morphology. (A) Normal appearance of a wing from a heterozygote Yorkie null (yki^B5^/+) fly. (B) A non-targeting RNAi control (gfp dsRNA) does not alter the wing morphology. RNAi knockdown of *CycE* or *Diap1* in the wing disc pouch using *nub*-GAL4 also does not disrupt wing morphology.

**Figure S4.**
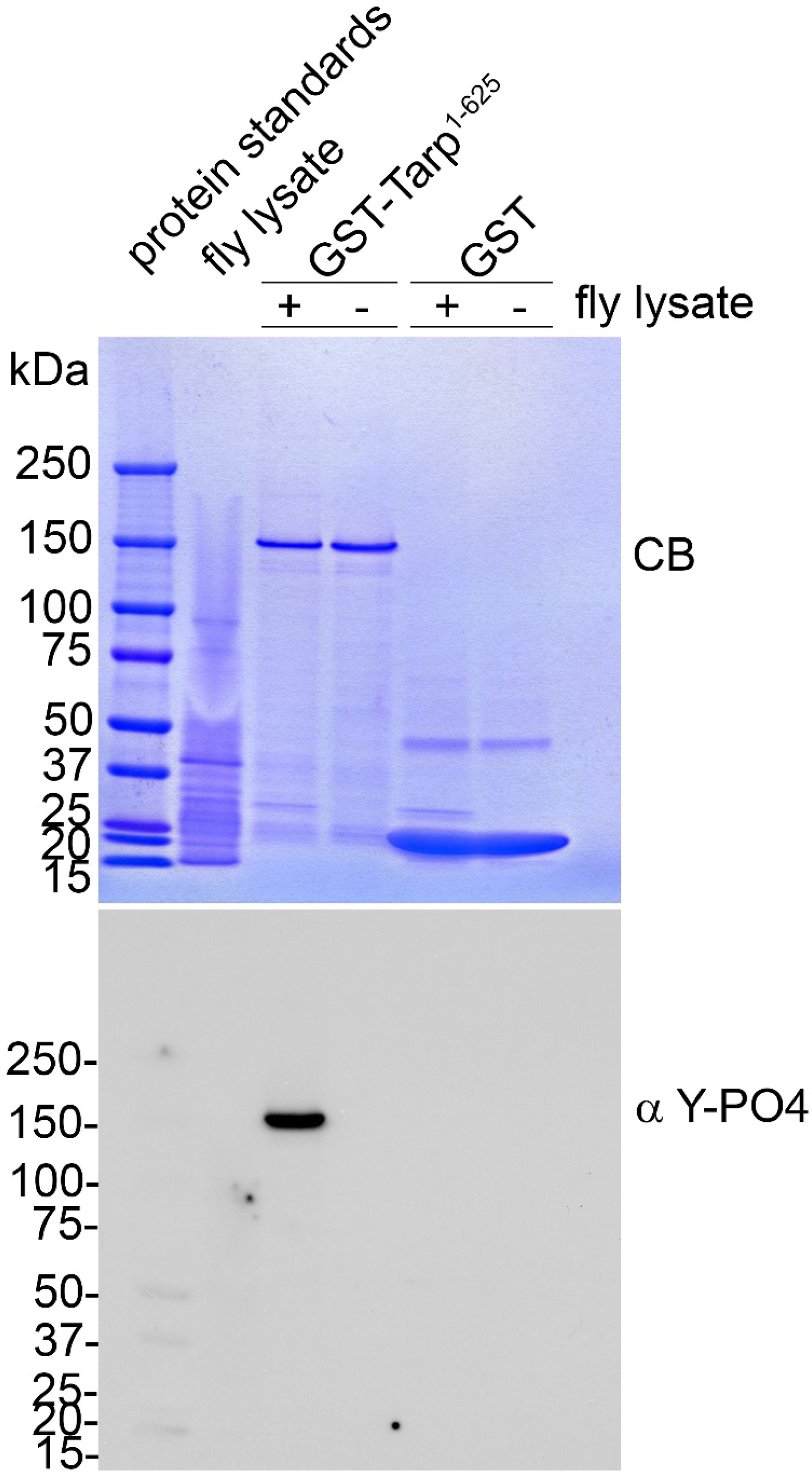
Endogenous *Drosophila* kinases can phosphorylate N-Tarp. (Top panel) Coomassie stain (CB) following SDS-PAGE validating the presence and relative amounts of the indicated samples. (Bottom panel) Western blot analysis of an identically loaded SDS-PAGE gel testing for the presence of tyrosine phosphorylation (α Y-PO4). Purified N-Tarp (GST-Tarp^1-625^) is tyrosine phosphorylated upon incubation with *Drosophila* lysate.

